# Investigating pathogenicity and virulence of *Staphylococcus pettenkoferi:* an emerging pathogen

**DOI:** 10.1101/2021.10.28.466297

**Authors:** Lucile Plumet, Nour Ahmad-Mansour, Sylvaine Huc-Brandt, Chloé Magnan, Alex Yahiaoui-Martinez, Karima Kissa, Alix Pantel, Jean-Philippe Lavigne, Virginie Molle

## Abstract

*Staphylococcus pettenkoferi* is a coagulase-negative *Staphylococcus* identified in 2002 that has been implicated in human diseases as an opportunistic pathogenic bacterium. Its multiresistant character is becoming a major health problem, yet the pathogenicity of *S. pettenkoferi* is poorly characterized. In this study, pathogenicity of a *S. pettenkoferi* clinical isolate from diabetic foot osteomyelitis was compared to a *Staphylococcus aureus* strain in various *in vitro* and *in vivo* experiments. Growth kinetics were compared against *S. aureus* and bacteria survival was assessed in the RAW 264.7 murine macrophage cell line, the THP-1 human leukemia monocytic cell line and the HaCaT human keratinocyte cell line. *Ex vivo* analysis were performed in whole blood survival assays, and *in vivo* assays via the infection model of zebrafish embryos. Moreover, whole-genome analysis was performed. Our results showed that *S. pettenkoferi* was able to survive in human blood, human keratinocytes, murine macrophages, and human macrophages. *S. pettenkoferi* demonstrated its virulence by causing substantial embryo mortality in the zebrafish model. Genomic analysis revealed virulence factors such as biofilm- (e.g., *icaABCD; rsbUVW*) and regulator- (e.g., *agr, mgrA, sarA, saeS*) encoding genes well characterized in *S. aureus*. This study thus advances the knowledge of this under investigated pathogen and validates the zebrafish infection model for this bacterium.

## Introduction

The genus *Staphylococcus* includes more than 50 species of Gram-positive cocci commonly classified into two categories in human medicine: coagulase-positive staphylococci (CoPS), mainly represented by *Staphylococcus aureus*, and coagulase-negative staphylococci (CoNS). CoNS are a diverse group of bacteria that range from true non-pathogenic to facultative pathogenic species with variable virulence potential. They share common characteristics involved in the transition to a pathogenic interaction with the host ^1^. As CoNS possesses less virulence factors than *S. aureus*, particularly those that cause infection, they are thought to be less harmful. However, CoNS are considered as opportunistic bacteria that colonize healthy people, as well as being the cause of the most common hospital infections with a growing effect on human health and life ^2^. Endogenous infections, such as bacteremia, endocarditis, osteomyelitis, pyoarthritis, peritonitis, mediastinitis, prostatitis, and urinary tract infections, are caused by CoNS present on the host’s skin and mucous membranes ^2^. These infections have been linked to over a dozen of the 50 CoNS species that have been identified, including *Staphylococcus capitis, Staphylococcus chromogenes, Staphylococcus cohnii, Staphylococcus epidermidis, Staphylococcus haemolyticus, Staphylococcus hominis, Staphylococcus lentus, Staphylococcus lugdunensis, Staphylococcus saprophyticus, Staphylococcus sciuri, Staphylococcus simulans, Staphylococcus warneri*, and *Staphylococcus xylosus* ^3^. Nevertheless, other rarer species have been reported, such as *Staphylococcus pettenkoferi* ^4^. Moreover, the therapeutic and prophylactic overuse of antibiotics contributes to the increase in infections due to multidrug-resistant CoNS ^2^.

*S. pettenkoferi* is a comparatively recently identified CoNS, for which its natural habitat, clinical implications, detection, and epidemiology remains to be characterized ^5^. Trülzsch *et al*., who first identified and described this pathogen in Germany in 2002, used 16S ribosomal ribonucleic acid (rRNA) sequencing, biochemical, and physiologic approaches to distinguish it from other CoNS ^4^. S. pettenkoferi bacteremia was initially described in a 25-year-old tuberculosis patient with extrapulmonary tuberculosis ^6^. Since then, around ten case reports of *S. pettenkoferi* bacteremia and one case of osteomyelitis from various countries have been published ^6–9^. Moreover, some *S. pettenkoferi* isolates have been detected as multidrug resistant to commonly used antibiotics such as beta-lactams, fluoroquinolones, and macrolides, complicating the treatment options ^10^.

Diabetic foot ulcers (DFU) represent one of the most significant complications for patients living with diabetes *mellitus*. These chronic wounds are responsible for frequent infections spreading to soft tissues and bone structures constituting diabetic foot osteomyelitis (DFOM), leading to increased amputations, mortality and morbidity ^11^. Whilst the infected DFU and DFOM are mainly polymicrobial ^12^, Grampositive cocci particularly *S. aureus* and CoNS are the most frequent bacteria isolated ^13^. Recently, we have started to observe an emergence of *S. pettenkoferi* isolated from bone biopsies in DFOM ^14^. Despite *S. pettenkoferi* being increasingly isolated in human pathology, its pathogenicity remains unexplored. Therefore, in order to evaluate the role of this bacterium as a pathogen, more exploratory investigations are needed. In this study, we demonstrate for the first time that *S. pettenkoferi* can persist in both macrophages and non-professionnal phagocytes, as well as in human blood, and its virulence was determined in the zebrafish model of infection.

## Materials and Methods

### Bacterial Strains, Media, and Growth Conditions

The bacterial strains used in this study are listed in Table 1. *S. pettenkoferi* SP165 was isolated from bone biopsies in a DFOM present in a 57-year-old man with Type-2 diabetes *mellitus* in the Gard-Occitanie Diabetic Foot Clinic (University hospital Nîmes, France). *Staphylococcus* strains were plated on Tryptic Soy Agar (TSA), or grown in Tryptic Soy Broth (TSB) medium at 37°C and 225 rpm. A microplate reader (Tecan, Trading AG) was used to monitor bacterial growth in 96-well plates.

**Table 1.** Strains used in this study

| Strain | Description | Resistance profile | Reference |
| --- | --- | --- | --- |
| SA564 | *S. aureus* clinical isolate, wild type | PEN | <sup>15</sup> |
| *S. pettenkoferi* SP165 | *S. pettenkoferi* clinical isolate from diabetic foot osteomyelitis, wild type | PEN, OXA, ERY, LIN, OFX, RIF, FOS | This study |

PEN, penicillin G; OXA, oxacillin; ERY, erythromycin; LIN, lincomycin; OFX, ofloxacin; RIF, rifampicin; FOS, fosfomicin

### Macrophage culture and infection

The murine macrophage cell line RAW 264.7 (mouse leukemic monocyte macrophage, ATCC TIB-71) was grown in Dulbecco’s Modified Eagle’s Medium (DMEM) (ThermoFisher Scientific) enriched with 10% fetal calf serum (ThermoFisher Scientific) at 37°C in a humidified environment with 5% CO2. The human leukemia monocytic cell line THP-1 ^16^ was cultured in Milieu Roswell Park Memorial Institute (RPMI) medium (ThermoFisher Scientific) enriched with 10% fetal calf serum (ThermoFisher Scientific) in a humidified environment at 5% CO2 at 37°C and differentiated into macrophages as previously described ^17^. *S. aureus* SA564 and *S. pettenkoferi* SP165 strains were cultured in TSB medium to the mid-exponential growth phase (OD_600_ = 0.7-0.9) for macrophage infection. The bacteria were collected for 5 minutes at 4000 rpm, rinsed in sterile PBS, centrifuged at 10000 rpm for 4 minutes, and finally resuspended in sterile PBS. THP-1 cells (1×10^6^ cells/mL, in 24 well-plates) and RAW 264.7 cells (5×10^5^ cells/mL, in 24 well-plates) were infected with *S. aureus* SA564 or *S. pettenkoferi* SP165 at a MOI of 20:1 (bacteria:cells) and incubated for 1 hour at 37°C and 5% CO2. After that, the cells were washed in PBS once more, and the extracellular bacteria were killed by incubation with gentamicin (100 μg/mL) for 30 minutes. After gentamicin treatment, macrophages were rinsed twice with PBS (T0), then incubated for 5 hours and 24 hours in fresh medium with 15 μg/mL lysostaphin for *S. pettenkoferi* SP165 and 5 μg/mL lysostaphin for *S. aureus* SA564. By lysing infected macrophages with 0.1 % Triton X-100 in PBS, intracellular bacteria were counted. To estimate the number of colony forming units(CFU), macrophage lysates were serially diluted and plated on TSB agar plates and grown at 37°C.

### Keratinocyte culture and infection

HaCaT, a well-known human keratinocyte cell line, is a spontaneously transformed human epithelial cell line from adult skin that retains complete epidermal differentiation potential ^18^. HaCaT cells were cultured in DMEM (ThermoFisher Scientific) enriched with 10% fetal calf serum (ThermoFisher Scientific), 1x Glutamate (Gibco ™), and 0.5% Penicillin/ Streptomycin antibiotics (Gibco ™) in a humidified environment at 37°C and 5% CO2. *S. aureus* SA564 and *S. pettenkoferi* SP165 were cultured for infection in TSB medium to the mid-exponential growth phase (OD_600_ = 0.7-0.9). The bacteria were then collected and resuspended in sterile PBS after centrifugation at 10,000 rpm for 5 minutes. The HaCaT cells (1×10^6^ cells/mL, in 24 well-plates) were infected with *S. aureus* SA564 or *S. pettenkoferi* SP165 at the MOI of 100:1 (bacteria:cells) and incubated 1h 30 at 37°C and 5% CO2. After that, the cells were washed in PBS once more, and the extracellular bacteria were killed by incubation with gentamicin (100 μg/mL) for 60 minutes. Following gentamicin treatment, macrophages were washed twice with PBS (T0), and then incubated for 5, 24, and 48 h in fresh medium with 15 μg/mL lysostaphin. By lysing infected HaCaT cells with 0.1 % Triton X-100 in PBS, intracellular bacteria were counted. Keratinocyte lysates were serially diluted and plated on TSB agar plates, and cultured at 37°C. The number of bacterial colonies at time post-gentamicin treatment (pGt) / number of bacterial colonies at T0 x 100 percent was used to calculate the survival rate of bacteria.

### Whole blood survival assay

*S. pettenkoferi* SP165 or *S. aureus* SA564 were incubated in freshly drawn blood, using EDTA as an anticoagulant. *S. aureus* SA564 and *S. pettenkoferi* SP165 strains were grown to the mid-exponential growth phase (OD_600_ = 0.7-0.9) in TSB. The bacteria were then resuspended in Roswell Park Memorial Institute medium (RPMI, Gibco) after centrifugation at 10,000 rpm for 5 minutes. Fresh venous human whole blood was obtained from healthy adult volunteers using EDTA-containing Vacutainer tubes (BD) (Etablissement Français du Sang, Montpellier, France) and inoculated with *S. aureus* SA564 or *S. pettenkoferi* SP165 at a concentration of 5×10^6^ bacteria/mL and incubated for 3 h at 37°C on a rotating wheel. The samples were serially diluted and plated on TSB agar plates, followed by incubation at 37°C. (CFUtimepoint/CFUinitialinput)*100 was used to compute the proportion of bacteria that survived.

### Infection of Danio rerio embryos

*S. aureus* SA564 and *S. pettenkoferi* SP165 strains were grown overnight at 37°C in TSB. Cultures were diluted 1:20 and grown to the mid-exponential growth phase (OD_600_ = 0.7-0.9) in TSB medium. The bacteria were then centrifuged for 10 minutes at 4000 rpm and reconstituted in fish water with 60 μg/mL sea salt (Instant Ocean) in distilled water with 4.10^-4^ N NaOH at a concentration of 2×10^7^ bacteria/mL. The number of bacteria in the inoculum was assessed by plating onto TSB agar plates after dilution in PBS. Experiments were carried out in fish water at 28°C with the GAB zebrafish line. Bath immersion infections were performed on embryos dechorionated at 48 hpf. Healthy embryos were put in groups of 24 in a Petri dish with the appropriate bacterial suspension (or fish water as a control) and then distributed individually into 96-well (Falcon) plates containing 200 μl of bacterial suspension (or fish water). Injured embryos were put on a Petri plate, anesthetized with 40 μg/mL tricaine, and a little transection of the tail fin was conducted under a stereomicroscope (Motic) with a 26-gauge needle to injure the tail fin prior infection. Following tail transection, groups of 24 injured embryos were put in a Petri dish with the appropriate bacterial suspension (or fish water as a control) and then dispersed individually into 96-well plates (Falcon) containing 200 μl of bacterial suspension (or fish water). The plates were held at a constant temperature of 28°C during the incubation process (bacteria are kept throughout the experiment in fish water, which does not support *S. aureus* SA564 or *S. pettenkoferi* SP165 growth). The absence of a heartbeat allowed the number of dead embryos to be counted visually.

### Ethical Statement

All animal experiments were carried out at the University of Montpellier in accordance with European Union recommendations for the care and use of laboratory animals (http://ec.europa.eu/environment/chemicals/labanimals/homeen.htm (accessed on 15 March 2021)) and were authorised by the Direction Sanitaire et Vétérinaire de l’Hérault and Comité d’Ethique pour l’Expérimentation Animale under reference CEEALR-B4-172-37 and APAFIS#5737-2016061511212601 v3. Adult zebrafish were not killed for this study, and breeding of adult fish followed the international norms set out by the EU Animal Protection Directive 2010/63/EU. According to the EU Animal Protection Directive 2010/63/EU, all studies were conducted prior to the embryos’ free feeding stage and did not constitute animal experimentation. The cardiac rhythm was used as a clinical criterion for survival graphs. Before bleach treatment, embryos were killed using the anesthetic tricaine at a fatal dosage (500 mg/mL).

### Whole-genome analysis

*S. pettenkoferi* strain SP165 was cultivated aerobically at 37°C for 24 h on Columbia sheep blood agar plates (5%) (Becton Dickinson). Following the manufacturer’s instructions, genomic DNA was extracted using the DNeasy UltraClean Microbial Kit (QIAGEN). Whole Genome Sequencing (WGS) was carried out using an Illumina MiSeq sequencing equipment (Illumina) using paired-end (PE) read libraries (PE250) made with an Illumina Nextera XT DNA Library Prep Kit (Illumina) according to the manufacturer’s instructions. To examine data quality, raw readings were processed using FastQC (v.0.11.9). To eliminate leftover PCR primers and filter low quality bases (Q_score <30) and short reads (<150bp), the Cutadapter tool (v.1.16) was used, which was implemented in Python (v.3.5.2). The downstream analysis includes the filtered trimmed reads. Using the CLC genomics workbench 7 (Qiagen), reads were mapped to the *S. pettenkoferi* FDAAGOS_288 genome (GenBank accession number: GCA_002208805.2) using default settings; length fraction: 0.5, similarity fraction: 0.8. Shovill v1.1.0 software on the Galaxy platform was used to process the assembled contigs for microbial genome annotation. The CONTIGuator website was used to produce consensus sequences, utilizing *S. pettenkoferi* FDAARGOS_288 as the reference. The virulence factor database (VFDB) was utilized to identify virulence factor-encoding genes from genome sequences (https://www.mgc.ac.cn/VFs/) ^19^. On assembled genomes, antimicrobial resistance genes were acquired from ABRIcate using the ResFinder database ^20,21^. The genome was annotated using PROKKA v1.14.5. The origin of replication was determined using GenSkew software (http://genskew.csb.univie.ac.at/) and plasmids were screened using the PlasmidFinder 2.0 webtool (https://cge.cbs.dtu.dk/services/PlasmidFinder/). WGS was subjected to other investigations, such as circular genome representation (using the BLAST Ring Image Generator program) (BRIG) ^22^). The newly sequenced strain can be found under the BioProject: PRJNA768314 for the raw reads and assembled genomes: SP165 SAMN22027592.

### Statistical Analyses

GraphPad software package Prism 6.01 was used to estimate the statistical significance of differences across groups, which is indicated in the relevant figure legends.

## Results

### S. pettenkoferi growth kinetics

*S. pettenkoferi* SP165 and *S. aureus* SA564 grown *in vitro* followed a typical growth cycle (Figure 1). A significant difference in exponential growth phase was observable between the two strains (p <0.01) in the first 8 h of growth, with *S. pettenkoferi* SP165 growing slower than *S. aureus*. The corresponding generation time (G = ln2 / μmax) was significantly different between *S. pettenkoferi* SP165 and *S. aureus* SA564: 74 min vs. 48 min for *S. aureus* SA564 (p<0.01). Furthermore, the differing growth patterns of the two isolates were represented in their colony size at 24 h on TSB agar plates, with *S. pettenkoferi* SP165 having a smaller colony size than *S. aureus* SA564 (data not shown).

**Figure 1.**
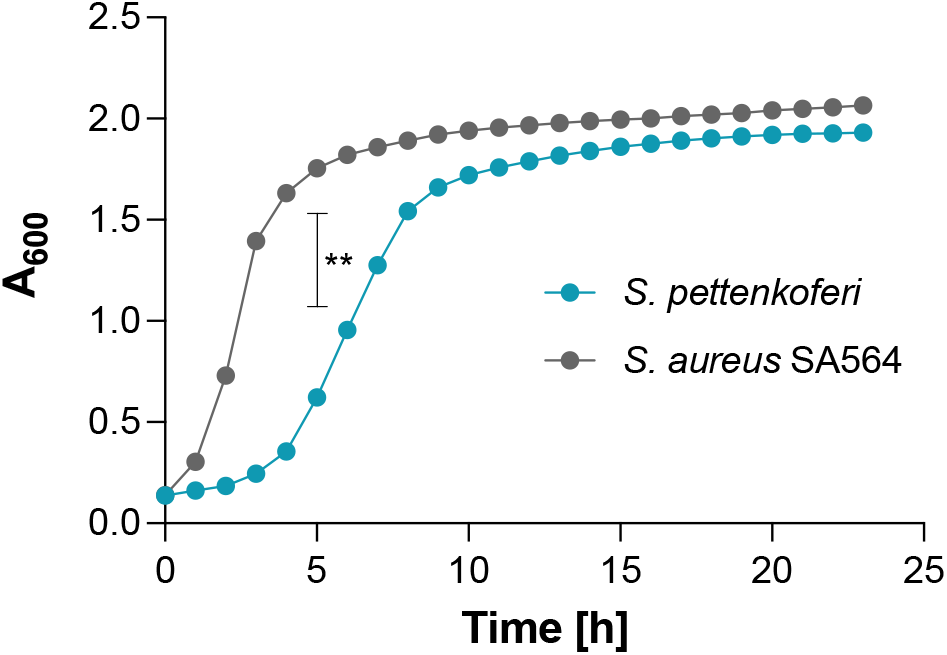
Growth kinetics of *S. pettenkoferi* SP165 (blue circles) and *S. aureus* SA564 (grey circles) strains in TSB. At a culture to flask volume of 1:10, cells were grown at 37°C and 225 rpm. At each time point (n = 3), the data show the mean A600 readings SD. Welsh’s t test, **, p0.01.

### S. pettenkoferi persists within murine and human macrophages

Macrophage immune cells have a key role in the eradication of bacteria via phagocytosis ^23,24^. To explore the interaction of *S. pettenkoferi* with macrophages, infection assays were conducted in the murine macrophage cell line RAW 264.7 and primary human THP-1 macrophages. Macrophages were allowed to phagocytose *S. pettenkoferi* SP165 or *S. aureus* SA564, and viable bacteria were recovered at 5 h and 24 h pGt. *S. aureus* SA564 and *S. pettenkoferi* SP165 were able to persist in both macrophage cell lines over time (Figure 2). Interestingly, intracellular *S. aureus* SA564 persisted without replication while *S. pettenkoferi* SP165 was able to replicate within 24 h pGt in RAW 264.7 macrophages (Figure 2A). Our data showed that both macrophage cell types failed to kill intracellular *S. pettenkoferi* SP165, but only human THP-1 cells restricted *S. pettenkoferi* SP165 bacterial growth (Figure 2B). These findings suggest that, in comparison to *S. aureus, S. pettenkoferi* SP165 has a distinct intracellular fate in which the bacteria can survive for long periods of time within macrophages. Therefore, the failure of macrophages to eradicate intracellular *S. pettenkoferi* may be a significant deficiency of host innate immunity, allowing for the existence of intracellular reservoirs of viable *S. pettenkoferi*.

**Figure 2.**
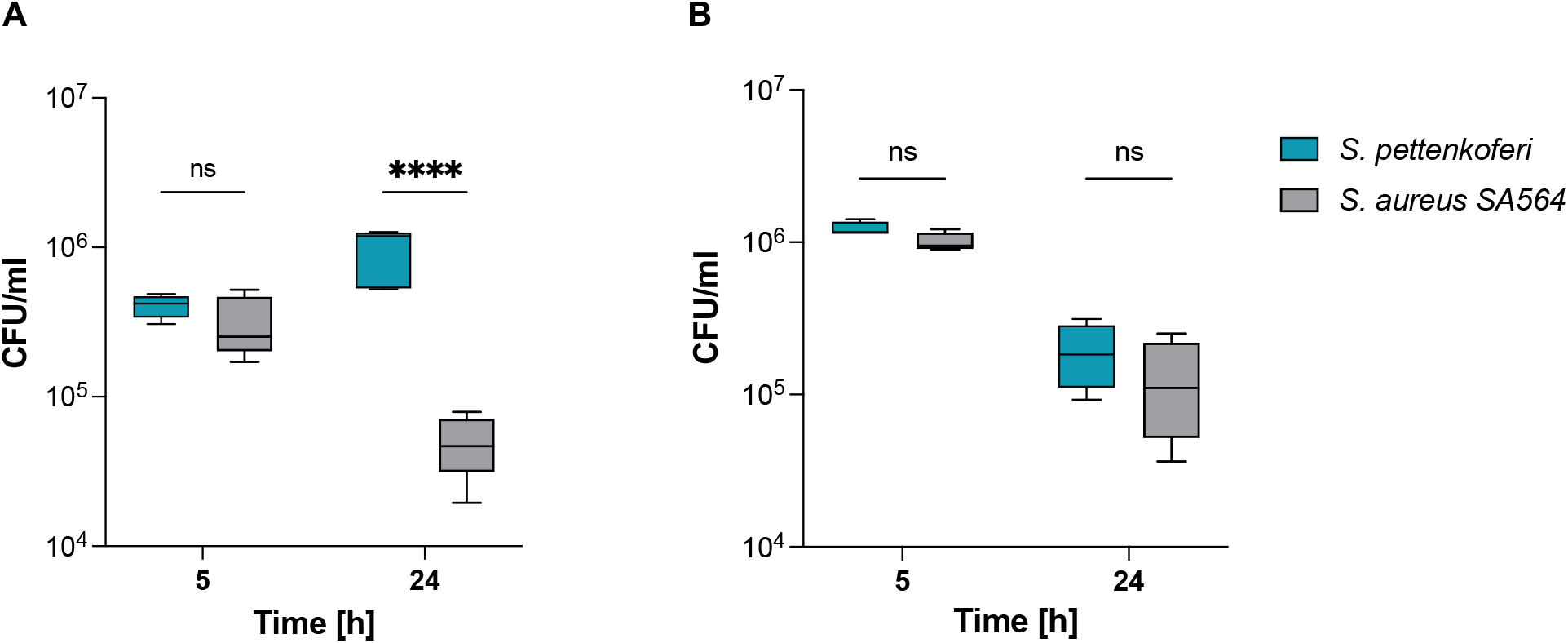
*S. pettenkoferi* SP165 survival in infected macrophages. *S. pettenkoferi* SP165 (blue) and *S. aureus* SA564 (grey) bacteria were used to infect RAW 264.7 (A) or THP-1 (B) macrophages. The amount of bacteria after 5 and 24 h pGt was evaluated after infection at a multiplicity of infection (MOI) of 20. The average and standard deviation (SD) of five different experiments are represented. A twoway ANOVA test was used to establish statistical significance, with ****, p<0.0001; ns, not significant

### S. pettenkoferi persists in human keratinocytes

*S. pettenkoferi* was identified in a case of osteomyelitis in a diabetic foot infection, indicating that this strain appears capable of infecting and persisting in non-phagocytic cells like osteoblasts or epithelial cells ^14^. Invasion of non-phagocytic host cells by *S. pettenkoferi* could be an effective mechanism to prevent elimination and maintain infection ^2^. To test this hypothesis, human skin keratinocytes were used to evaluate the survival capacity of *S. pettenkoferi* SP165. HaCaT is a non-transformed human keratinocyte line that has been used to investigate epithelial infection ^25^, cytotoxicity ^26^, biofilm formation ^27^, and skin cancer ^28^. It can also be employed in models of skin wound infection where the stratum corneum is disrupted and bacteria come into close contact with living keratinocytes ^27^. Therefore, we performed invasion experiments to address the persistence of *S. pettenkoferi* SP165 since *S. aureus* has been demonstrated to proliferate and persist intracellularly in HaCaT cells ^29,30^. A significant reduction of *S. aureus* SA564 intracellular bacteria compared to *S. pettenkoferi* SP165 was observed over 48h in HaCaT cells (Figure 3).

**Figure 3.**
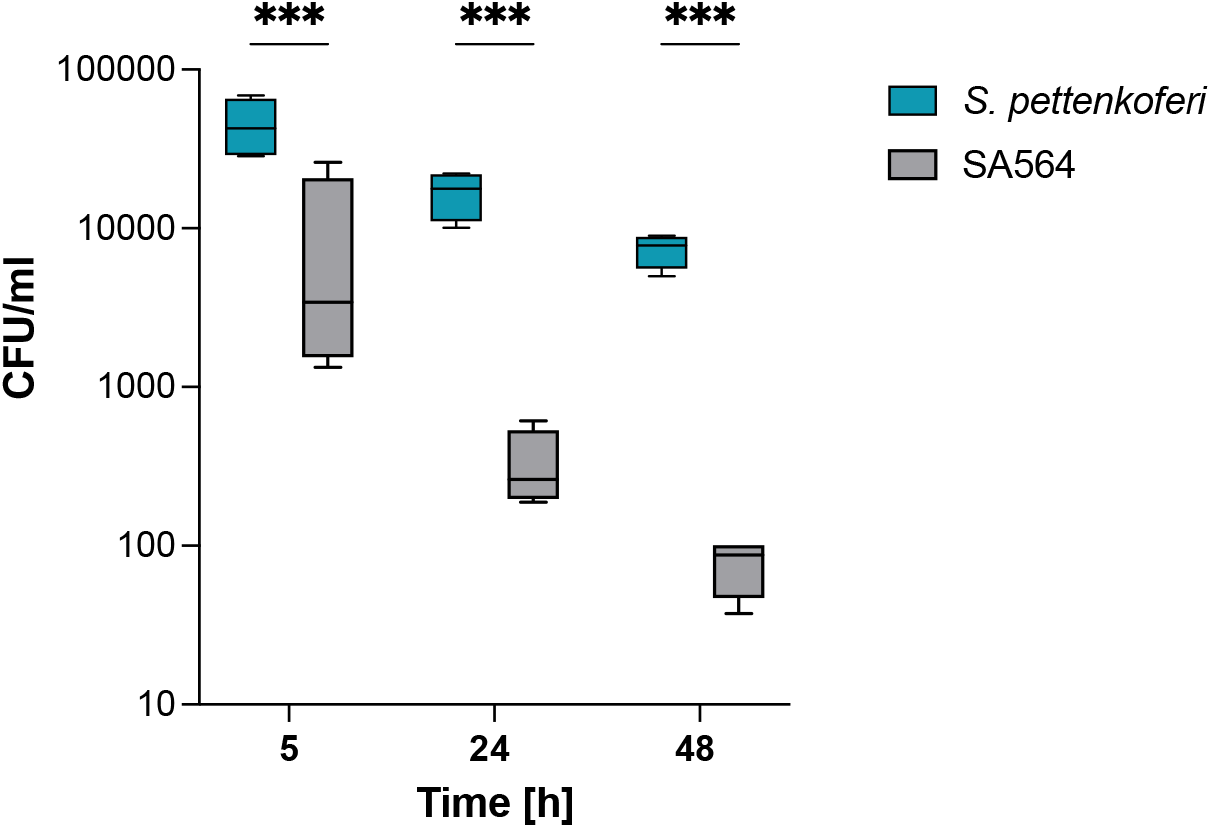
Survival capacity of *S. pettenkoferi* SP165 in non-professional phagocytes. At a MOI of 100, bacteria from *S. pettenkoferi* SP165 (blue) and *S. aureus* SA564 (grey) were utilized to infect cells of the keratinocyte cell line HaCaT, and the cells were co-cultured for 90 minutes. Washing and lysostaphin/gentamicin treatment were used to eliminate extracellular and adhering bacteria, and infected cells were grown for up to 48 h in cell culture media supplemented with lysostaphin. Eukaryotic cells were lysed, and surviving bacteria in lysates were evaluated by counting CFUs at 5, 24, and 48 h after lysostaphin/gentamicin treatment. Percent survival = (#CFUfinal/#CFUinput)*100. Statistical significance was determined by two-way ANOVA test where ***, p<0.001.

### S. pettenkoferi avoids whole blood killing

*Ex vivo* survival of *S. pettenkoferi* in human blood was evaluated to define the pathogen’s capacity to induce disseminated infection. After 3 h of incubation in human blood, 100% of the *S. pettenkoferi* SP165 inoculum survived, but only 10% of the *S. aureus* SA564 inoculum survived (p<0.05). (Figure 4). Interestingly, despite a slower growth rate for *S. pettenkoferi* SP165 in early growth (Figure 1), *S. pettenkoferi* SP165 showed around a 10-fold higher rate of survival in blood than *S. aureus* SA564.

**Figure 4.**
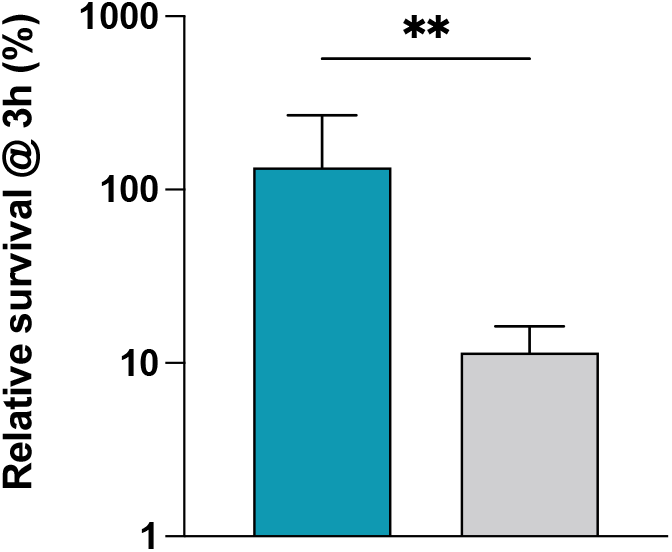
*S. pettenkoferi* SP165 and *S. aureus* SA564 survival in whole blood. Percent survival of *S. pettenkoferi* SP165 (blue) and *S. aureus* SA564 (grey) in freshly collected human blood. Bacteria were cultivated overnight, diluted, inoculated in blood, maintained for 3 h at 37°C, and rotated. Percent survival = (CFU_timepoint_/CFU_initialinput_)*100. The data is the average with standard deviation of five separate experiments. **, p<0.01 (Mann-Whitney U test).

### A bath infection model using healthy and wounded zebrafish embryos was used to evaluate S. pettenkoferi pathogenicity

The zebrafish (Danio rerio) is being studied for its ability to mimic human diseases caused by bacterial pathogens ^31^. It is a common model for studying hostpathogen interactions ^32^. Microinjecting *S. aureus* bacteria into the bloodstream of 1- or 2-day postfertilization (dpf) old embryos is a frequent method to generate *S. aureus* infections ^33^. Acute infection and death result when the number of bacteria injected exceeds the embryo’s phagocytic capability. To avoid the time-consuming microinjection step, we employed a robust and efficient bath immersion model for *S. pettenkoferi* infection in zebrafish embryos. This innovative approach, which uses wounded embryos to measure bacterial pathogenicity, has already been validated for other pathogens ^34,35^. Two separate models were employed for bath immersion infections: immersion alone and immersion following injury, in which the animals’ tail fin tips were wounded prior to infection. The same bacterial solution at the same concentration was used to infect healthy and wounded larvae in a parallel experiments. Bath immersion was first performed on healthy embryos at 2 days post fertilization, while the mouth was still closed ^36^. The viability of zebrafish embryos was monitored for 48 h, and no deaths were recorded with *S. aureus* SA564, while less than 20% of embryos survived 24 h post infection (hpi) following *S. pettenkoferi* SP1165 infections (Figure 5A). In injured larvae infected with *S. pettenkoferi* SP165, mortalities began at 18 hpi and increased up to 100% by 30 hpi, while embryos infected with *S. aureus* SA564 showed a mortality around 30% by the end of the assay (Figure 5B). There were no mortality among healthy or wounded larvae treated with fish water at the end of the studies (control group). Therefore, our results confirmed the use of this model to study *S. pettenkoferi* virulence.

**Figure 5.**
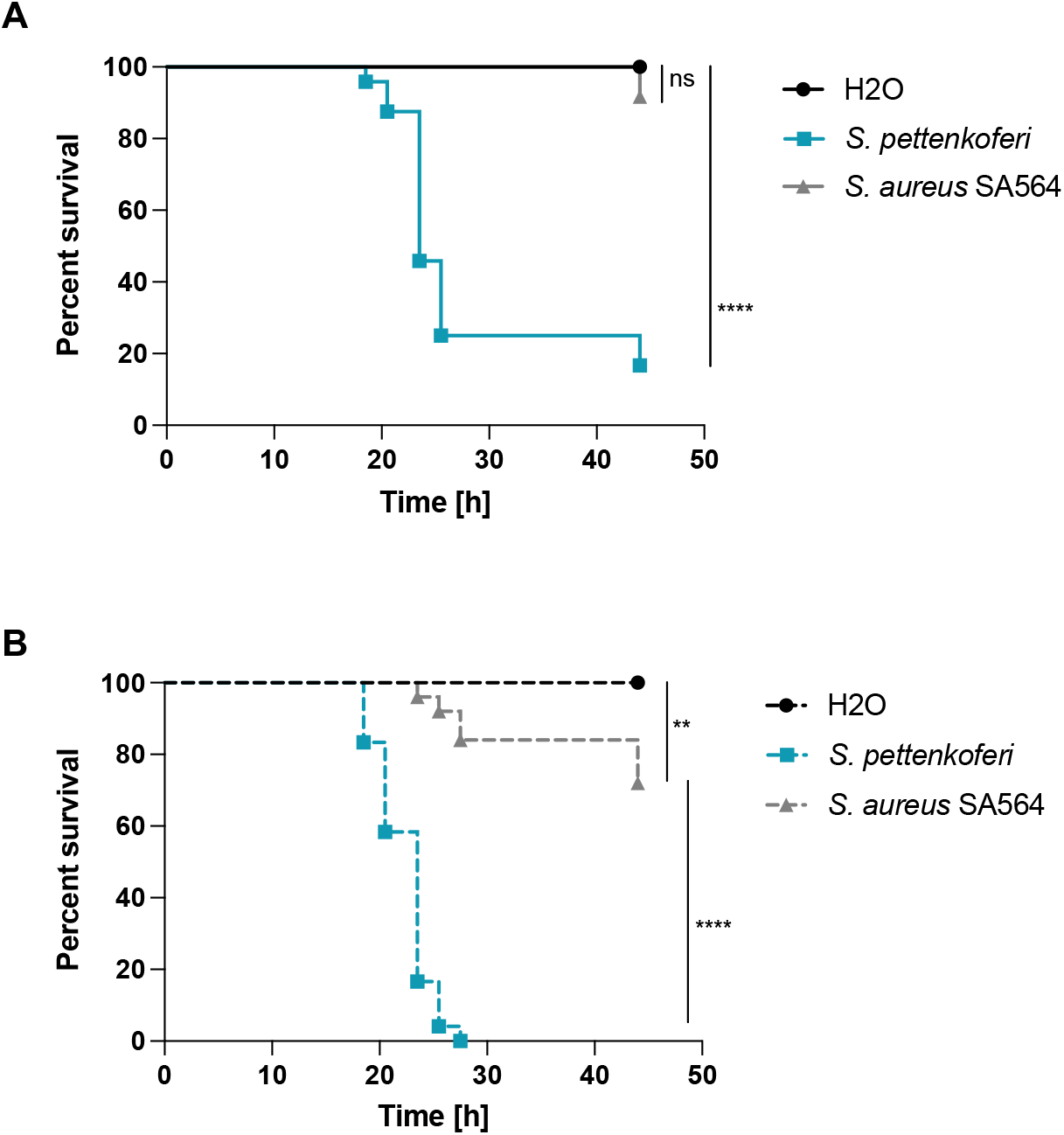
Virulence of *S. pettenkoferi* SP165 and *S. aureus* SA564 strains in zebrafish embryos. Kaplan-Meier representation of the survival of **(A)** healthy zebrafish embryos or **(B)** zebrafish embryos injured in the tail fin at 48 h post-fertilization (hpf) in a bath infected with *S. pettenkoferi* SP165 or *S. aureus* SA564 strains at 2.10^7^ CFU/mL grown in exponential phase, or “fish water” (negative control). The proportion of surviving embryos (n = 24 for each, indicative of three separate experiments) is used to express the results. Significant difference at ** p<0.01, **** p<0.0001 or no significant difference (ns).

### Whole-genome analysis of S. pettenkoferi SP165

The virulence potential of SP165 was investigated by whole genome sequencing. The genome size of *S. pettenkoferi* genomes corresponded to 2,435,720 base pairs (bp), a smaller genome size compared to the two other sequenced strains (Table 2). The *S. pettenkoferi* strains exhibited a similar GC content with 38.91%, 38.85% and 39.14% for SP165, FDAARGOS 288 and FDAARGOS 1071, respectively. The numbers of coding DNA sequences (CDS) were 2,298 for SP165 compared to 2,380 for FDAARGOS 288 and 2,318 for FDAARGOS 1071. BRIG-based analyses are shown in Figure 6. The origin of replication was estimated at 1,409,866 bp. The genomic comparison between the *S. pettenkoferi* SP165 and the two strains, FDAARGOS 288 and FDAARGOS 1071, showed a good coverage between each genome with a high identity (90% coverage and 96.75 and 99.92% identity, respectively). Different virulence factors were identified detecting some well-known biofilm- (e.g., *icaABCD; rsbUVW*) or regulator- (e.g., *agr, mgrA, sarA, saeS*) encoding genes present in *S. aureus* (Table S1). For resistome analysis, the detected mechanisms of resistance corroborated the antibiogram results with: methicillinoresistance (*mecA*), fluoroquinolones resistance (mutations in *gyrA* and *gyrB*), macrolides resistance (*msrAB*), rifampicin resistance (*rpoB, rpoC*), and fosfomycin resistance (*fosB*) markers.

**Table 2.** General characteristics of *S. pettenkoferi* SP165 strain.

| Features | SP165 | FDAARGOS 288 | FDAARGOS 1071 |
| --- | --- | --- | --- |
| Biosample | SAMN22027592 | SAMN06173301 | SAMN16357240 |
| Assembly | SHOVILL 1.1.0 | CA v. 8.2 | SMRT v. 7.1.0, HGAP v. 4 |
| Genome size (bp) | 2,435,720 | 2,502,360 | 2,449,395 |
| Number contigs | 54 | ND | ND |
| GC Content (%) | 38.91 | 38.85 | 39.14 |
| Number CDS | 2,298 | 2,380 | 2,318 |
| Year of collection | 2021 | 2014 | ND |
| Year of sequencing | 2021 | 2021 | 2020 |
| Locality | Nîmes (France) | Washington (USA) | Braunschweig (Germany) |

ND, not determined; CDS, coding DNA sequences

**Figure 6.**
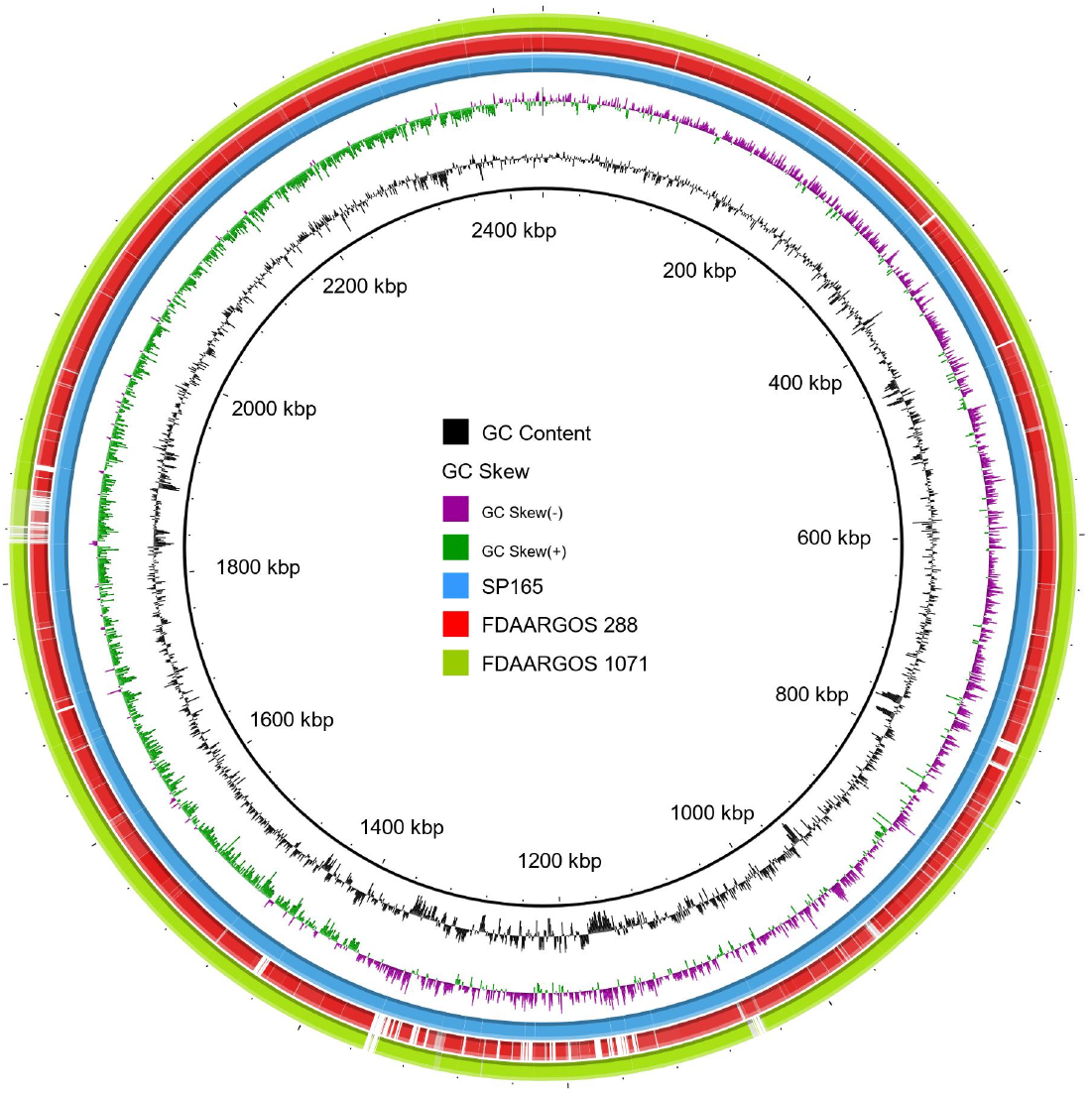
BRIG analysis of *S. pettenkoferi* genomes. The SP165 strain was compared against the two genomes previously described in the literature. The inner blue ring represents the SP165 genome; the middle red one shows the FDAARGOS 288 genome after BLASTn match; while the outer green ring corresponds to FDAARGOS 1071. Only regions with > 90% nucleotide identity are colored. Lower identity percentage or no match are represented by blank spaces in each ring. The inner bicolor ring corresponds to the GC skew of *S. pettenkoferi* SP165 isolates. Green profile indicates overabundance of GC nucleotides whereas purple shows the opposite. The inner black ring represents the GC%.

## Discussion

*S. pettenkoferi* was first identified in 2002 ^4^, and is part of the normal skin microbiota; however, this bacterium can cause infection, with several cases of severe infection reported ^6–9^. However, to date, *S. pettenkoferi* pathogenic processes and virulence have not been described. Here, we report that *S. pettenkoferi* SP165 isolated from DFOM is a slow-growing species that can either persist in both macrophages and non-professional phagocytes, as well as in human blood. Moreover, its virulence, investigated for the first time in the zebrafish model of infection, was clearly demonstrated.

In this study, *S. pettenkoferi* SP165 was characterized by a slower exponential growth phase in planktonic growth correlated to a late colony appearance time. Bacterial fitness is generally measured by the rate at which bacterial cells multiply when grown in well-aerated, nutrient-rich medium. Bacteria potential to outcompete their opponents by rapidly expanding must be evaluated against the threat of depleting available nutrients, meaning a prolonged time of starvation. Faster-growing cells, on the other hand, displayed not only a larger number than their rivals, but they might be more resistant against antibiotics according to Gutierrez et *al*. results. New experimental evolution investigations and a detailed examination of bacterial survival mechanisms have helped researchers to figure out if pathogen growth is connected to pathogenicity ^37^. Numerous mathematical models have been proposed during the last 30 years to characterize the link between rate of growth and virulence heterogeneity ^38^. One of the basic ideas in such models is that it exists a link between growth rate and pathogenicity, with greater growth rates enhancing dissemination but also promoting host mortality. However, Legget *et al*. showed that slow-growing pathogens were a lot more virulent than fast-growing pathogens, and that pathogens breathed or infecting via cutaneous lesions were far more virulent than those swallowed ^38^. More recently, poor correlations between virulence measures and replication rates suggest that, in addition to growth rate, other characteristics could account for the variations in virulence, like the potential to control the host immune system ^39^. Therefore, the relatively slow growth rate of *S. pettenkoferi* SP165 compared to *S. aureus* SA564 could be an advantage to persist during host infection.

Furthermore, we established that *S. pettenkoferi* SP165 may survive and proliferate in professional phagocytes as well as in murine macrophages. The death of bacteria upon phagocytosis is due to the collaborative effort of immunological factors, involving NADPH oxidase stimulation and the generation of reactive oxygen radicals ^40,41^, acidification of phagosomes ^42^, peptidoglycan hydrolysis mediated by lysozymes, and activation of lysosomal proteases ^23^. Despite this, previous *in vitro* and *in vivo* studies have demonstrated that *S. aureus* may survive in the phagolysosomes of macrophages, where it begins multiplying ^43–46^. Recently, it was demonstrated that *S. lugdunensis*, a CoNS bacterium that can cause significant infection, has the ability to bypass host immunity. Flannagan *et al*. showed that ingested *S. lugdunensis* are not killed by macrophages, and that the bacteria can survive in mature phagolysosomes for long periods without multiplying ^46^. Conversely, *S. pettenkoferi* intracellular persistence might represent a survival strategy to prevent intoxication of the host cell. In the absence of alternative evading processes, this might hypothetically enable internalized *S. pettenkoferi* to avoid extrinsic immunological factors, that could result in bacterial killing. The murine and human macrophages were not able to eradicate phagocytosed *S. pettenkoferi*, showing that this pathogen is able to evade the antibacterial properties of phagolysosomes, whereas the mechanisms remain unknown. Furthermore, *S. pettenkoferi* SP165’s failure to grow inside human macrophages varies markedly from that of murine immune cells, highlighting the functional distinctions between RAW 264.7 and primary human macrophages. RAW264.7 macrophages lack a functional caspase-1 inflammasome activity ^47,48^, potentially contributing to this replication difference, as this pathway is critical for bacterial clearance despite *S. aureus* having developed many strategies to regulate cell death processes such as apoptosis, necroptosis, and pyroptosis in order to generate infection ^49^. Furthermore, *S. pettenkoferi*’s ability to survive inside macrophages could be critical in systemic infections in which Kupffer cells remove staphylococci from blood ^45,50^. Kupffer cells rapidly remove *S. aureus* from the bloodstream and, due to their inability to eradicate phagocytosed *S. aureus*, provide this pathogen with a secure intracellular habitat in which to grow ^45,51^. Although it remains to be determined whether *S. pettenkoferi* can survive in Kupffer cells. Moreover, we demonstrated *S. pettenkoferi* SP165’s ability to persist in non-professional phagocytes such as human keratinocytes without replicating or being outwardly toxic to the host cells. Therefore, our data support the notion that macrophages or keratinocytes capacity to limit *S. pettenkoferi* intracellular growth is undeniably anti-bacterial; yet, the inability to eliminate ingested cocci is a weakness that might enable *S. pettenkoferi* to remain within the human body, using macrophages as bacterial reservoirs. However, the mechanisms involved in intracellular survival of *S. pettenkoferi* remain to be established.

CoNS including *S. pettenkoferi* are becoming more widely recognized as a primary cause of bacteremia, particularly in patients with medical implants, such as intravenous catheters, artificial heart valves, and joint prosthetics, or those who are immunodeficient ^6,52^. In the case of *S. aureus*, its ability to thrive in the human body is dependent on a delicate balance between its various virulence factors and the presence of different host defense mechanisms. *S. aureus* can breach the epithelial and endothelial barriers to reach the bloodstream, regardless of the original location of infection ^53,54^. The bacteria come into contact with the innate immune system in the blood, which is made up primarily of neutrophils, monocytes, and the complement system. In this study, the antimicrobial properties of whole human blood were examined towards *S. pettenkoferi* SP165 and *S. aureus* SA564 strains, revealing that *S. pettenkoferi* possesses an approximately 10-fold higher rate of survival than *S. aureus* SA564. Surprisingly, while CoNS do not possess the coagulase that is considered as an important virulence factor for blood survival, they are often associated with bacteremia ^2^. However, coagulase binds to host prothrombin and forms staphylothrombin, which activates thrombin protease activity. Regardless of the fact that coagulase is considered to help protect bacteria from phagocytic and immunological responses by causing localised coagulation, its involvement in pathogenicity is unknown ^55^. In the case of the *S. epidermidis* CoNS, the induction of coagulase expression dramatically reduced its survival in the blood, suggesting that coagulase synthesis in the blood could be unfavorable to survival ^55^. This contradicts the theory that coagulase protects bacteria from immune defenses by producing localized coagulation. It remains to be determined whether the extraordinary capacity of *S. pettenkoferi* to survive in whole human blood is linked to the absence of the coagulase or other mechanisms such as the ability to evade immune clearance as seen in *S. epidermidis* ^56,57^. In addition, the genome of this bacterium revealed high homology with *S. aureus* concerning the presence of virulence factors and the main regulators of this staphylococcal virulence. Their role in the pathogenicity of *S. pettenkoferi* must be elucidated.

In addition, to examine *S. pettenkoferi* pathogenicity, we employed a newly established procedure for infection of zebrafish embryos. The zebrafish model offers a number of benefits compared to mammalian infection models related to technical, economic, and ethical issues. Zebrafish are vertebrates, which are genetically and physiologically closer to humans than invertebrate models, and have a functioning innate immune system in embryos ^58^. Because both macrophages and neutrophils play a role in preventing *S. aureus* development, zebrafish have improved our comprehension of *S. aureus* mechanisms developped to avoid the host innate immunity ^59^, but some can operate as an immune bottleneck, preserving a subpopulation of bacteria from being killed by host cells and causing disseminated infection ^60,61^. Interestingly, zebrafish infection studies demonstrated how interactions with commensals on human skin might increase *S. aureus* colonization ^62^. In most cases, bacterial infections in zebrafish embryos are performed by microinjection ^35^. *S. pettenkoferi* is a bacterium that enters via skin wounds ^2^, thus we applied a bath infection model that mimics real infection. We showed that when healthy or wounded embryos were submerged in *S. pettenkoferi* bacteria at 2 dpf (a developmental time when the mouth is not yet open), significant mortality was reported, demonstrating *S. pettenkoferi* virulence. Surprisingly, even at 2 dpi, *S. pettenkoferi* is virulent, indicating that without mouth ingestion, this pathogen is able to induce mortality in contrario to *S. aureus* SA564. Fin excision has previously been used as a model of “sterile” wounding damage and inflammation ^63^. An inflammatory response is triggered by the recrutment of neutrophils and macrophages following an injury, and the embryonic zebrafish fin is also remarkably regenerative ^64^. Therefore, *S. pettenkoferi* seems able to manipulate the immune host response to avoid killing in this infection model. Interestingly, while zebrafish cannot replace other vertebrate models like mice; they can disclose key principles in *S. pettenkoferi* virulence and host defense, and thus support the development of new treatments for staphylococcal diseases.

## Acknowledgments

We acknowledge Sarah Kabani for her help with the editing. We thank the structural, human, and financial assistance provided by the Nîmes University Hospital via the award “Thématiques phares.” The researchers are members of the FHU InCh (Federation Hospitalo Universitaire Infections Chroniques, Aviesan).

## Data availability statement (DAS)

On reasonable request, the corresponding author will provide the datasets produced during and analysed in the present work.

## Transparency declaration

All the authors report no conflicts of interest relevant to this editorial.

## Author Contributions

Conceptualization, J.P.L, V.M.; investigation, L.P., N.A.M, S.H-B, C.M.; resources, K.K., A.Y-M.; writing–original draft preparation, L.P, J.P.L. and V.M.; writing—review and editing, K.K.., S.H-B, C.M.; visualization, N.A.M., S.H-B.; supervision, J.P.L and V.M.; project administration, J.P.L and V.M.; funding acquisition, J.P.L and V.M. All authors have read and agreed to the published version of the manuscript.

