## Supplementary Table S1 for "Investigating pathogenicity and virulence of *Staphylococcus pettenkoferi:* an emerging pathogen"

**Supplementary data**

**Table S1.** Main resistome and virulome genes present in *S. pettenkoferi* SP165.

| Gene | Function |
|---|---|
| **Resistome** | |
| *fosB* | Fosfomycin |
| *gyrA* | Fluoroquinolone |
| *gyrB* | Fluoroquinolone |
| *katA* | Nitric oxide |
| *mecA* | Oxacillin |
| *msrC* | Erythromycin |
| *parC* | Fluoroquinolone |
| *parE* | Fluoroquinolone |
| *rpoB* | Rifampin |
| *rpoC* | Rifampin |
| *ugpQ* | Associated with mecA |
| *msrAB* | Macrolide efflux pump |
| *qacC* | Quaternary ammonium compound |
| *sdrM* | Norfloxacin, acriflavine and ethidium bromide |
| *phnB* | Bleomycin |
| *copC/copD* | Copper |
| *hicB_lk_antitox* | Nutrient deprivation, antibiotic treatment, and immune system attacks |
| **Virulome** | |
| *agrA* | *agr* complex |
| *agrB* | *agr* complex |
| *agrC* | *agr* complex |
| *agrD* | *agr* complex |
| *cvfB* | Increases expression of virulence factor |
| *gapA1* | Glyceraldehyde-3-phosphate dehydrogenase 1 |
| *gapA2* | Glyceraldehyde-3-phosphate dehydrogenase 2 |
| *icaA* | Biofilm production |
| *icaB* | Biofilm production |
| *iIcaC* | Biofilm production |
| *icaD* | Biofilm production |
| *lyrA* | Lysostaphin resistance protein A |
| *lss* | Bacteriocin against *S. aureus* |
| *mgrA* | Regulator of *agr* |
| *rsbU* | Biofilm production |
| *rsbV* | Biofilm production |
| *rsbW* | Biofilm production |
| *saeS* | Regulation of staphylococcal virulence factors |
| *sarA* | Regulator of *agr* |
| *sarR* | Regulator of *agr* |
| *sarX* | Regulator of *agr* |
| *sarZ* | Regulator of *agr* |

| | |
|---|---|
| *setC* | Detoxification of non-metabolizable sugar analogs |
| *sigB* | Biofilm production |
| *traP* | Regulator of *agr* |
| *vraS* | Biofilm production |
| *isaB* | Expressed during septicemia |
| *mprF* | Resistance mechanism against cationic antimicrobial peptides (CAMP) |
| *brxA*/*brxB* | Resistance to infection by bacteriophage VR7 and VpaE1 |
| *pulG* | Nutrient acquisition (TSS II pathway) |
| *yihY* | Resistance to complement-dependent killing by serum |
| *liaF* | Cell wall-active antibiotics response protein |
| DUF4097 | Putative adhesin |
| *rhbC* | LucA/LucC family siderophore biosynthesis protein |
| IDKPNINI_00836 | Putative holin-like toxin |
| *yaaT* | Sporulation |
| IDKPNINI_02349 | Virulence factor |
